# CDE-1 suppresses the production of risiRNA by coupling polyuridylation and degradation of 26S rRNA

**DOI:** 10.1101/2019.12.18.880609

**Authors:** Yun Wang, Chenchun Weng, Xiangyang Chen, Xufei Zhou, Xinya Huang, Meng-Qiu Dong, Chengming Zhu, Shouhong Guang

**Author notes:** These authors contributed equally to this work (Yun Wang, Chenchun Weng).

## Abstract

Antisense ribosomal siRNAs (risiRNAs) downregulate pre-rRNAs through the nuclear RNAi pathway in *Caenorhabditis elegans*. However, the biogenesis and regulation of risiRNAs remain obscure. Previously, we showed that 26S rRNAs are uridylated at the 3’-ends by an unknown terminal polyuridylation polymerase before the rRNAs are degraded by a 3’ to 5’ exoribonuclease SUSI-1(ceDIS3L2). There are three polyuridylation polymerases, CDE-1, PUP-2, and PUP-3, in *C. elegans*. Here, we found that CDE-1 is specifically involved in suppressing risiRNA production. CDE-1 localizes to perinuclear granules in the germline and uridylates both Argonaute-associated 22G-RNAs and 26S rRNAs at the 3’-ends. Immunoprecipitation followed by mass spectrometry (IP-MS) revealed that CDE-1 interacts with SUSI-1(ceDIS3L2). Consistent with those results, both CDE-1 and SUSI-1(ceDIS3L2) are required for the inheritance of RNAi. Therefore, this work identified a rRNA surveillance machinery of rRNAs that couples terminal polyuridylation and degradation.

## Introduction

RNAs are extensively modified: 5’ termini are often capped, internal positions are altered on both ribose rings and bases, and 3’ termini receive untemplated nucleotides, which are referred to as tails. In eukaryotes, tails occur on most classes of RNAs, and they control RNA processing, stability, transport and function. Terminal modification is critical in biology. For example, uridylation is implicated in tumorigenesis, proliferation, stem cell maintenance, and immune defense against viruses and retrotransposons (Blahna, Jones et al., 2011, Hagan, Piskounova et al., 2009, Jones, Quinton et al., 2009, Le Pen, Jiang et al., 2018, Warkocki, Krawczyk et al., 2018, Yeo & Kim, 2018). The *C. elegans* genome encodes three polyuridylation polymerases (PUPs): *cde-1/pup-1/cid-1*, *pup-2* and *pup-3* (Kwak & Wickens, 2007). These PUPs may have distinct roles in different cellular contexts. *cde-1* is involved in the inheritance of RNAi, chromosome segregation and antiviral defense (Le Pen et al., 2018, van Wolfswinkel, Claycomb et al., 2009, Xu, Feng et al., 2018). CDE-1 functions with the RNA-dependent RNA polymerase (RdRP) EGO-1 and the Argonaute CSR-1 in the germline to affect chromosome segregation (Claycomb, Batista et al., 2009). PUP-2/3 are the homologues of TUT4/7 (terminal uridylyl transferases 4/7) in mammals. PUP-2 targets the microRNA *let-7* and regulates the stability of LIN-28 (Lehrbach, Armisen et al., 2009). The balance of CDE-1, PUP-2 and PUP-3 activities appears to ensure proper germline development in *C. elegans* (Li & Maine, 2018).

Ribosome biogenesis is a very sophisticated multistep process, in which mistakes can occur at any step. Cells must carefully surveil the steps of the pre-rRNA processing and the assembly of ribosomal subunits. Misprocessed rRNAs are usually surveyed and degraded by multiple supervision machineries, including the exosome complex and the Trf4/Air2/Mtr4p polyadenylation (TRAMP) complex, etc. (Henras, Plisson-Chastang et al., 2015, Lafontaine, 2010, Lafontaine, 2015). Aberrant RNAs are degraded by exosomes in a 3’-5’ exonucleolytic decay manner (Houseley, LaCava et al., 2006, Thoms, Thomson et al., 2015, Vanacova & Stefl, 2007). The exosome-independent exoribonuclease DIS3L2 plays a pivotal role in the 3’-5’ degradation of oligouridylated RNA fragments (Faehnle, Walleshauser et al., 2014, Lubas, Damgaard et al., 2013, Pirouz, Munafo et al., 2019, Ustianenko, Pasulka et al., 2016, Zhou, Feng et al., 2017b).

In addition to degrading erroneous rRNAs, antisense ribosomal siRNAs (risiRNAs) silence pre-rRNAs through the nuclear RNAi pathway to suppress the accumulation of erroneous rRNAs in *C. elegans* (Yan, Zhu et al., 2019, Zhou, Chen et al., 2017a, Zhou et al., 2017b, Zhu, Yan et al., 2018). Erroneous rRNAs are usually oligouridylated at the 3’-ends and then degraded by the exoribonuclease SUSI-1(ceDis3L2). However, it is unclear which terminal uridyltransferase performs the untemplated addition of the 3’-end uracil. Identifying which PUP is involved in the 3’-uridylation of erroneous rRNAs and how it is involved will further our understanding the quality control mechanism of cellular nucleic acids.

Here, we found that CDE-1 is specifically involved in suppressing risiRNA production. CDE-1 localizes to perinuclear granules in the germline and uridylates both Argonaute-associated 22G-RNAs and 26S rRNAs at the 3’-ends. Interestingly, we found that CDE-1 interacts with SUSI-1(ceDIS3L2). Both CDE-1 and SUSI-1(ceDIS3L2) are required for the inheritance of RNAi. Therefore, we conclude that CDE-1 suppresses the generation of risiRNAs by uridylating 26S rRNA and recruiting SUSI-1(ceDIS3L2) to the rRNA.

## Results

### Depletion of CDE-1 promotes risiRNA production

There are three RNA terminal uridylyltransferase genes, *cde-1, pup-2*, and *pup-3*, that are involved in RNA 3’ uridylation in *C. elegans.* We previously showed that risiRNA was enriched in WAGO-4-bound siRNAs in *cde-1* mutants (Xu et al., 2018). To further study the specificity and function of *cde-1* in risiRNA production, we used the *GFP::NRDE-3* transgene as a reporter. NRDE-3 is an Argonaute protein that transports siRNAs from the cytoplasm to the nucleus (Guang, Bochner et al., 2008). NRDE-3 localizes to the nucleus when it binds to siRNAs, but it accumulates in the cytoplasm when not bound to siRNA ligands. Disruption of the generation of endogenous siRNAs, for example, in the *eri-1* mutant result in relocalization of NRDE-3 from the nucleus to the cytoplasm. We crossed *eri-1(mg366);GFP::NRDE-3* onto the *pup* mutant lines and found that the depletion of *cde-1*, but not *pup-2* or *pup-3*, was able to redistribute NRDE-3 from the cytoplasm to the nucleus (Figure 1A). We generated a single copy transgene *CDE-1::mCherry* by MosSCI technology. This transgene was able to rescue the *cde-1(tm936)* defects and redistribute NRDE-3 from the nucleus to the cytoplasm (Figure S1A). To exclude the possibility that PUP-2 and PUP-3 act redundantly to suppress siRNA generation, we generated *pup-2;pup-3* double mutants. In the double mutants, NRDE-3 still accumulated in the cytoplasm (Figure S1B). These data suggest that NRDE-3 was bound to newly generated siRNAs in *cde-1* mutants.

**Figure 1.**
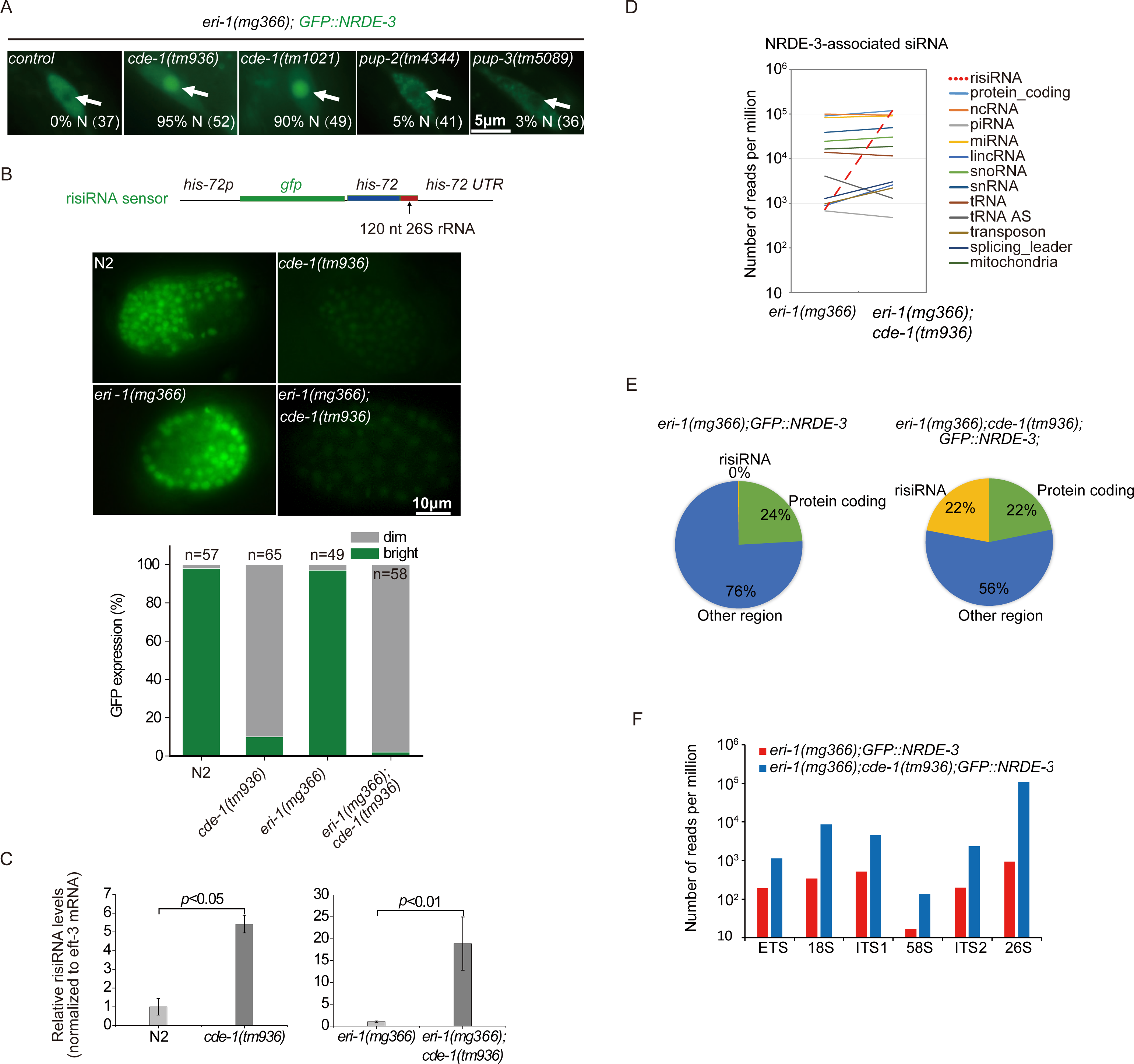
Antisense ribosomal siRNA (risiRNA) accumulated in *cde-1* mutants. (A) NRDE-3 localized to the nucleus in *eri-1(mg366);cde-1(tm936);GFP::NRDE-3* animals. Images show representative seam cells of *C. elegans*. The number of scored animals is indicated in the parentheses. White arrows, nucleus. (B) The risiRNA sensor is silenced in the *cde-1* mutant. Indicates are imagers of late embryos. The levels of GFP expression were scored in the bottom panel. (C) qRT-PCR analysis of risiRNA levels in indicated animals. Data are presented as the mean ± s.d. n = 3. (D-E) Deep sequencing of NRDE-3-associated siRNAs in indicated animals. The red dashed line indicates risiRNAs. (F) The number of risiRNAs targeting the each region of pre-rRNA transcription unit were analyzed.

To test whether the NRDE-3-bound siRNAs in *cde-1* mutants contain risiRNA sequences, we used a risiRNA sensor expressing a *his-72p::gfp::his-72* reporter fused to the 26S rRNA sequence (Figure 1B). The sensor was expressed in wild-type N2 and *eri-1(mg366)* animals but silenced in *cde-1(tm936)* mutants. We quantified the amount of risiRNA by quantitative real-time PCR analysis and found that risiRNAs were increased in *cde-1* mutants (Figure 1C).

Last, we immunoprecipitated NRDE-3 and deep sequenced its associated small RNAs in *eri-1(mg366);GFP::NRDE-3* and *eri-1(mg366);cde-1(tm936);GFP::NRDE-3* animals in a 5’-phosphate-independent manner. Notably, the proportion of NRDE-3-bound risiRNAs increased approximately 164-fold in *eri-1(mg366);cde-1(tm936);GFP::NRDE-3* animals compared to the values observed in control animals (Figures 1D and 1E). The abundance of risiRNAs targeting each rRNA region increased in *cde-1(tm936)* animals (Figure 1F).

To search for the genetic requirements of risiRNA production in the *cde-1* mutants, we crossed *rrf-1*, *rrf-2*, and *rrf-3*, lines onto the *eri-1(mg366);cde-1(tm936);GFP::NRDE-3* animals. RRF-1, RRF-2, and RRF-3 are RNA-dependent RNA polymerases that are important for the generation of 22G-RNAs in *C. elegans.* Consistent with previous results, the depletion of *rrf-1* and *rrf-2* together resulted in NRDE-3 being redistributed from the nucleus to the cytoplasm (Figure S1C). In addition, the depletion of *rrf-1* and *rrf-2* together partially restored the fecundity of *eri-1;cde-1* animals (Figure S1D).

We conclude that *cde-1* likely acts as a suppressor of siRNA (*susi)* gene and suppresses the generation of risiRNAs.

### CDE-1 uridylates risiRNA

We first compared the small RNA expression profiles between wild-type and *cde-1* mutant animals. Small RNAs were isolated from young adult animals through use of the TRIzol reagent and deep sequenced in a 5’-phophate-independent manner. Although the depletion of *cde-1* did not noticeably change the expression profile of different small RNA categories, risiRNAs were enriched 4.7 fold in *cde-1* mutant animals vs wild-type animals (Figure 2A). We then immunoprecipitated GFP::NRDE-3 and deep sequenced the associated siRNAs. NRDE-3-bound risiRNAs were enriched 17 fold in *cde-1* mutant animals, compared to what was observed in control animals (Figure 2B).

**Figure 2.**
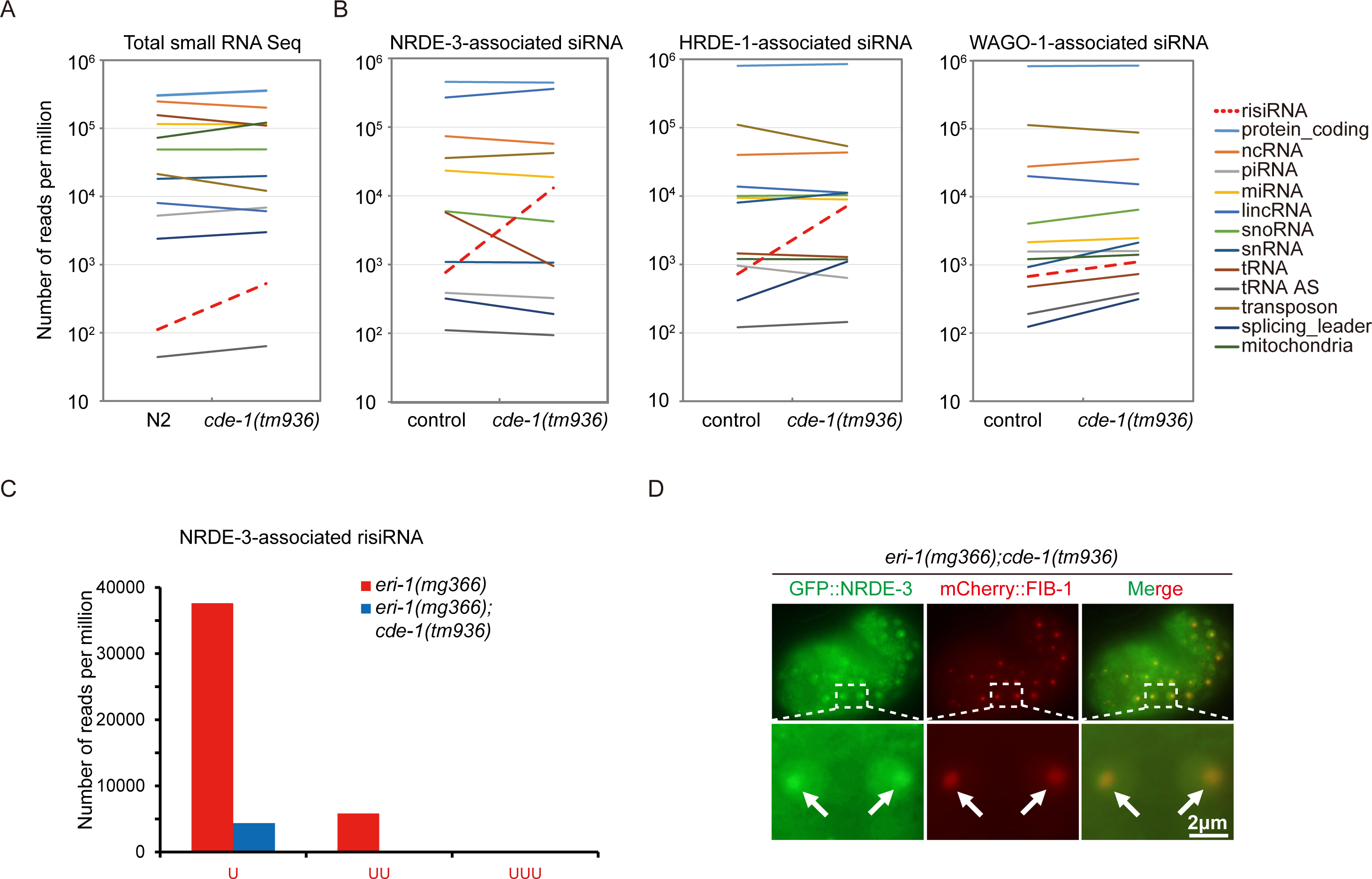
CDE-1 uridylated risiRNA. (A) Deep sequencing of total small RNAs of indicated animals. The red dashed line indicates risiRNAs. (B) Deep sequencing of Argonaute-associated siRNAs in indicated animals. (C) Number of uridylated NRDE-3-associated risiRNAs in indicated animals. (D) risiRNA elicited nucleolar accumulation of NRDE-3 in *cde-1* mutants. Indicated were images of late embryos of *eri-1(mg366);cde-1(tm936);GFP::NRDE-3;mCherry::FIB-1*. White arrows, nucleolus.

To test whether risiRNAs bind to Argonaute proteins in addition to NRDE-3, we analyzed HRDE-1 and WAGO-1-bound small RNAs in the young adult animals. HRDE-1 and WAGO-1 were immunoprecipitated from the control animals and the *cde-1(tm936)* mutant animals. Small RNAs were isolated and deep sequenced in the 5’-phosphate-independent method. In *cde-1* mutants, the amount of risiRNAs bound to HRDE-1 and WAGO-1 increased 9.9- and 1.6-fold, respectively, compared to those bound in wild-type animals (Figure 2B). The NRDE-3-, HRDE-1-, and WAGO-1-bound small RNAs still exhibited the characteristics of 22G-RNAs, which is 22 nt in length and starts with 5’ guanidine in the mutants (Figure S2). A similar increase in risiRNA was observed in WAGO-4-bound risiRNAs in *cde-1* mutants (Xu et al., 2018). CDE-1 adds untemplated uracil to the 3’-ends of CSR-1- and WAGO-4-bound siRNAs. We analyzed the NRDE-3-bound risiRNAs and found that there was a loss of the added untemplated uracil in *cde-1* mutants (Figure 2C).

Small RNAs associate with NRDE-3 and guide NRDE-3 to the target nuclear nucleic acids. In the presence of risiRNA, NRDE-3 accumulated in the nucleoli of *cde-1* mutants (Figure 2D). FIB-1 in *C. elegans* is encoded by an ortholog of the genes encoding human fibrillarin and *Saccharomyces cerevisiae* Nop1p (Lee, Lee et al., 2012, Yi, Ma et al., 2015). FIB-1 localizes to the nucleolus in embryos.

### CDE-1 interacts with SUSI-1(ceDIS3L2) in the germline

To further understand the function of CDE-1, we constructed a GFP::3*×*FLAG tagged *cde-1p::CDE-1::GFP::3×FLAG* transgene (abbreviated as *CDE-1::GFP*) using Mos1-mediated single-copy insertion (MosSCI) technology. CDE-1 was expressed in the germline cells at all developmental stages (Figure S3A). We noticed that CDE-1 accumulated in both the cytoplasm and the perinuclear region exhibiting distinct foci in the germline of adult animals. We crossed the *CDE-1::GFP* strain with the P-granule marker strain *mRuby::PGL-1*, and found that perinuclear localized CDE-1 largely colocalized with the P-granule marker PGL-1 (Figure 3A).

**Figure 3.**
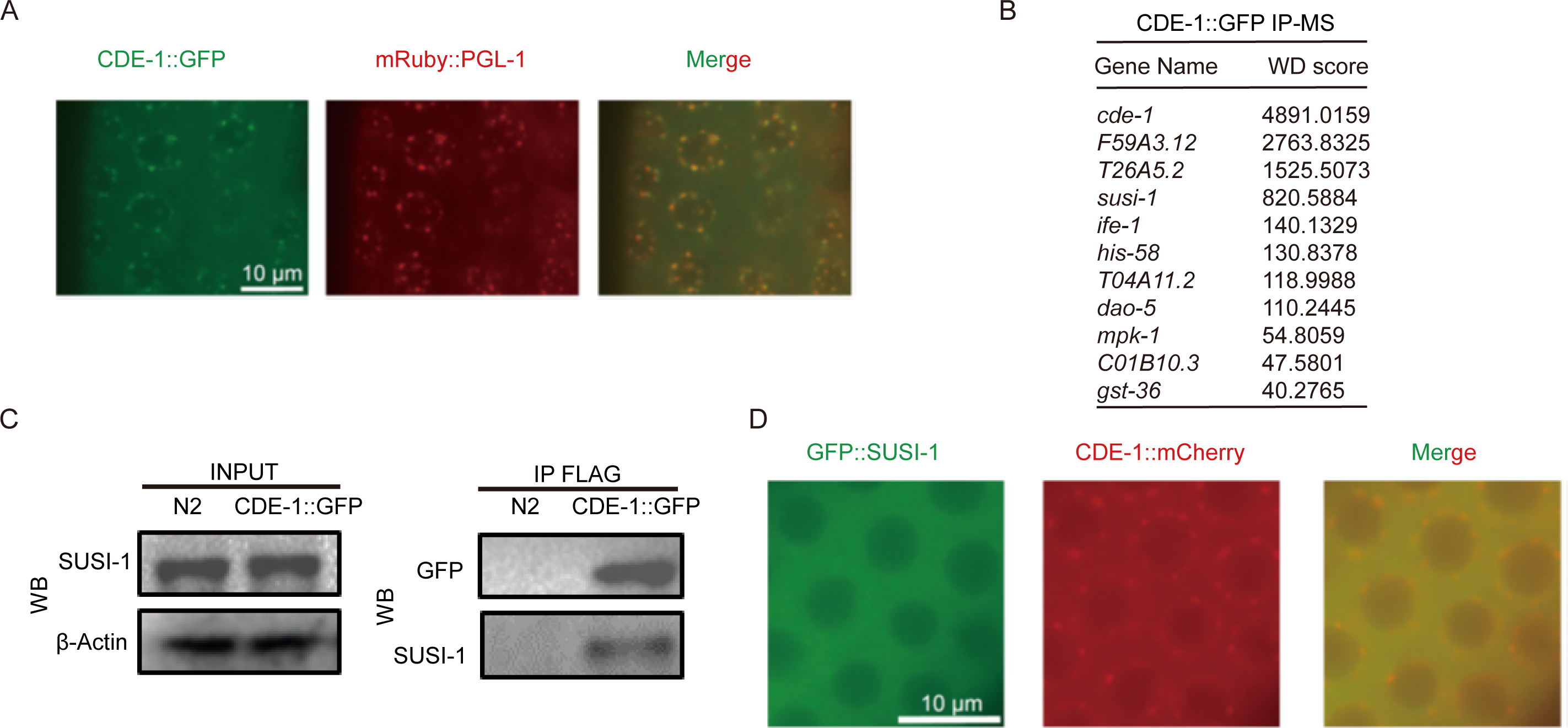
CDE-1 interacted with SUSI-1(ceDIS3L2) in the germline. (A) CDE-1 largely colocalized with the P-granule marker PGL-1. Images show the germline cells of young adult animals. (B) A summary of the top ten putative interacting proteins identified by CDE-1 immunoprecipitation followed by mass spectrometry. (C) The protein-protein interaction of CDE-1 and SUSI-1(ceDIS3L2) was assayed by coimmunoprecipitation followed by western blotting with the indicated antibodies. (D) SUSI-1(ceDIS3L2) accumulated in the cytoplasm. Indicates were germline cells of young adult animals.

We searched for proteins that interact with CDE-1. We used coimmunoprecipitation followed by mass spectrometry (IP-MS) to identify proteins that potentially interact with CDE-1. Strikingly, we identified SUSI-1(ceDis3L2) (Figure 3B and Figure S3B). SUSI-1 is a 3’ to 5’ exoribonuclease that degrades oligouridylated RNAs. In *susi-1* mutants, both risiRNAs and oligouridylated rRNAs accumulate (Zhou et al., 2017b). To confirm the protein-protein interaction between CDE-1 and SUSI-1, we generated an antibody targeting SUSI-1(ceDis3L2). CDE-1::GFP was immunoprecipitated by anti-FLAG antibody. Western blotting of the pelleted proteins with SUSI-1(ceDis3L2) antiserum confirmed the protein-protein interaction between CDE-1 and SUSI-1(ceDis3L2) *in vivo* (Figure 3C). We then generated single-copy *3×FLAG::GFP::SUSI-1* and *CDE-1::mCherry* transgenes and found that SUSI-1(ceDis3L2) accumulated in the cytoplasm of the germline (Figure 3D).

Therefore, we conclude that CDE-1 and SUSI-1 likely function as a protein complex to suppress risiRNA production.

### CDE-1 is involved in uridylation of 26S rRNA

Previously we showed that SUSi-1 degrades oligouridylated rRNAs and suppresses the production of risiRNA (Zhou et al., 2017b). To test whether CDE-1 uridylates rRNAs, we used a 3’ tail-seq assay to examine whether rRNA that was oligouridylated at 3’-tail was depleted in *cde-1(tm936)* animals (Figures 4A and 4B). Total RNA was isolated from embryos and L3-staged control and *cde-1* animals, ligated to a barcoded DNA linker and reverse transcribed with a primer complementary to the linker. Libraries were then prepared by PCR with a 26S rRNA primer and a primer targeting the linker. Illumina adaptor sequences were subsequently added, which was followed by a number of PCR cycles and high-throughput sequencing.

**Figure 4.**
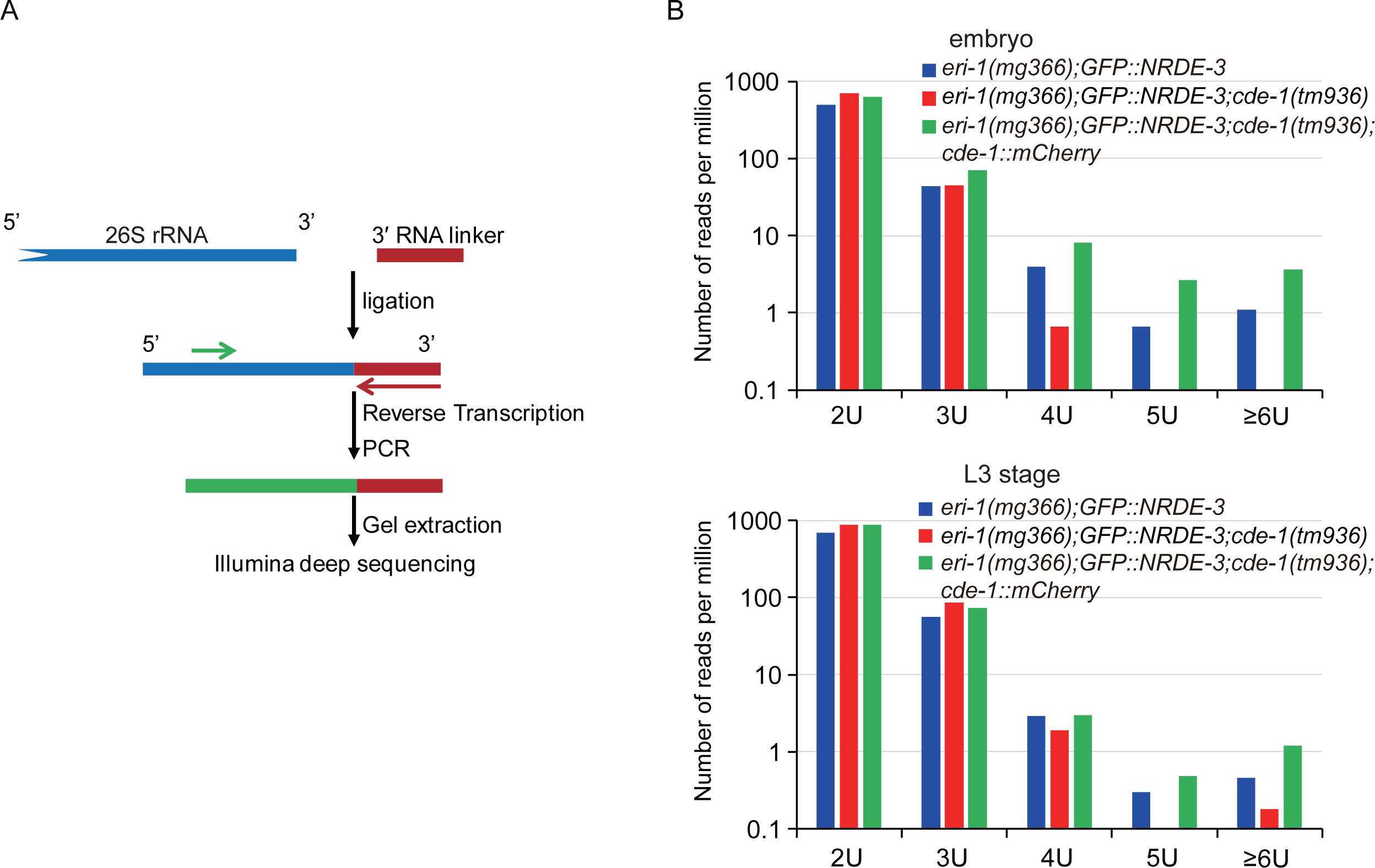
CDE-1 was involved in the 3’-end uridylation of 26S rRNAs. (A) A schematic of the rRNA tail-seq method. (B) Tail-seq data of 26S rRNAs from indicated animals at the embryo and L3 stages. The number of reads with untemplated oligouridylation at the 3’-ends is indicated.

The 3’-end of 26S rRNA was extensively modified by all four nucleotides compared to the annotated rRNA sequence (Zhou et al., 2017). Only a small fraction of the 3’-end exactly matched to the annotated 26S rRNA from the Wormbase WS250 assembly. Although we did not detect a dramatic change in the nontemplated addition of a single nucleotide, we observed a modest depletion of oligouridylation at the 3’-tail of 26S rRNA, comparing *cde-1(tm936)* to control animals (Figure 4B). The introduction of the *CDE-1::mCherry* transgene can rescue this oligouridylation defect. We conclude that CDE-1 is involved in uridylating rRNAs.

### SUSI-1(ceDIS3L2) is required for the inheritance of RNAi

It was previously showed that CDE-1 is required for the inheritance of RNAi by uridylating WAGO-4-associated siRNAs (Spracklin, Fields et al., 2017, Xu et al., 2018). Since CDE-1 interacts with SUSI-1(ceDIS3L2), we tested whether *susi-1* was also required for the inheritance of RNAi. We used a germline-expressed *mex-5p::GFP::H2B* (abbreviated as *GFP::H2B*) transgene as a reporter, which can inherit RNAi-induced gene silencing for multiple generations. Both *hrde-1* and *cde-1* were not required for exogenous *gfp* dsRNA to silence the *GFP::H2B* transgene in the parental generation, but they were essential for silencing in the F1 generation (Figures 5A and 5B). Similarly, *susi-1(ceDis3L2)* was not required for exogenous *gfp* dsRNA to silence the *GFP::H2B* transgene in the P0 generation, but was necessary for silencing in F1 progeny. We conclude that *susi-1(ceDis3L2)* is required for the inheritance of RNAi.

**Figure 5.**
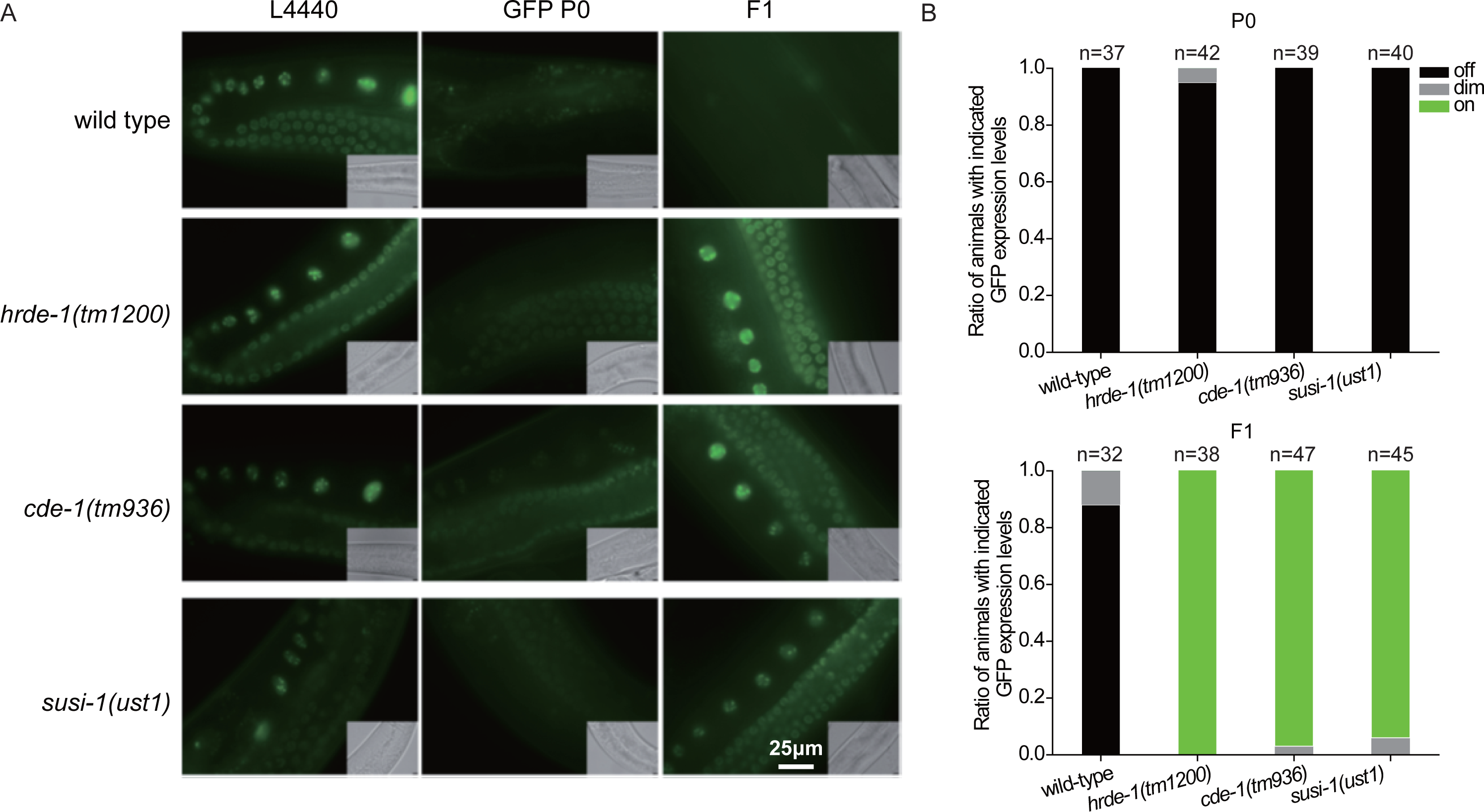
Both CDE-1 and SUSI-1(ceDIS3L2) were required for the inheritance of RNAi. (A) *mex-5p::GFP::H2B* transgenic animals were exposed to bacteria expressing *gfp* dsRNA. F1 embryos were isolated and grown on control bacteria in the absence of further *gfp* dsRNA treatment. GFP expression in the indicated animals was imaged in the germline and oocytes. (B) The percentage of P0 and F1 animals expressing GFP was counted.

## Discussion

Misprocessed rRNAs are usually detected and degraded by surveillance machinery during ribosome biogenesis. Previously, our lab identified a class of antisense ribosomal siRNAs (risiRNAs) that downregulate pre-rRNAs through the nuclear RNAi pathway to suppress the accumulation of erroneous rRNAs. We identified a number of broadly conserved genes that are involved in rRNA processing and maturation. The depletion of these genes lead to an increase in risiRNAs. Thereafter, these genes are named suppressor of siRNA (*susi*). Among them, SUSI-1(ceDIS3L2) plays a vital role in the 3’-5’ degradation of oligouridylated rRNA fragments. In this work, we further found that CDE-1 uridylates the 3’-end of 26S rRNAs and recruits SUSI-1(ceDIS3L2) through protein-protein interactions. Therefore, we conclude that *cde-1* is a new *susi* gene and suppresses the generation of risiRNAs.

Uridylation of the 3’-end of RNAs plays important functions in determining the fate of RNA (Lee, Kim et al., 2014, Menezes, Balzeau et al., 2018). For example, uridylation is an intrinsic step in the maturation of noncoding RNAs, including the U6 spliceosomal RNA or mitochondrial guide RNAs in trypanosomes (Trippe, Guschina et al., 2006). Uridylation can also switch specific miRNA precursors from a degradative to a processing mode. This switch depends on the number of uracils added and is regulated by the cellular context (De Almeida, Scheer et al., 2018, Heo, Ha et al., 2012). However, the typical consequence of uridylation is accelerating the RNA degradation (Pirouz, Du et al., 2016, Ustianenko et al., 2016). In this work, we showed that CDE-1 can uridylate 26S rRNAs and recruit the 3’-5’ exoribonuclease SUSI-1(ceDIS3L2), which may further promote the degradation of oligouridylated rRNAs. In the absence of either CDE-1 or SUSI-1(ceDIS3L2), erroneous rRNAs will accumulate in cells, which thereafter recruit the RNA-dependent RNA polymerases, including RRF-1 and RRF-2, to initiate risiRNA production (Figure 6). risiRNAs then bind to both nuclear and cytoplasmic Argonaute proteins and silence rRNAs through both nuclear and cytoplasmic RNAi machinery. Therefore, risiRNA and the RNAi machinery, together with exoribonucleases, act to avoid the accumulation of potentially harmful or unnecessary erroneous rRNA transcripts (Henras et al., 2015, Houseley et al., 2006, Karbstein, 2013, Lafontaine, 2010, Pena, Hurt et al., 2017, Schmidt & Butler, 2013, Vanacova & Stefl, 2007).

**Figure 6.**
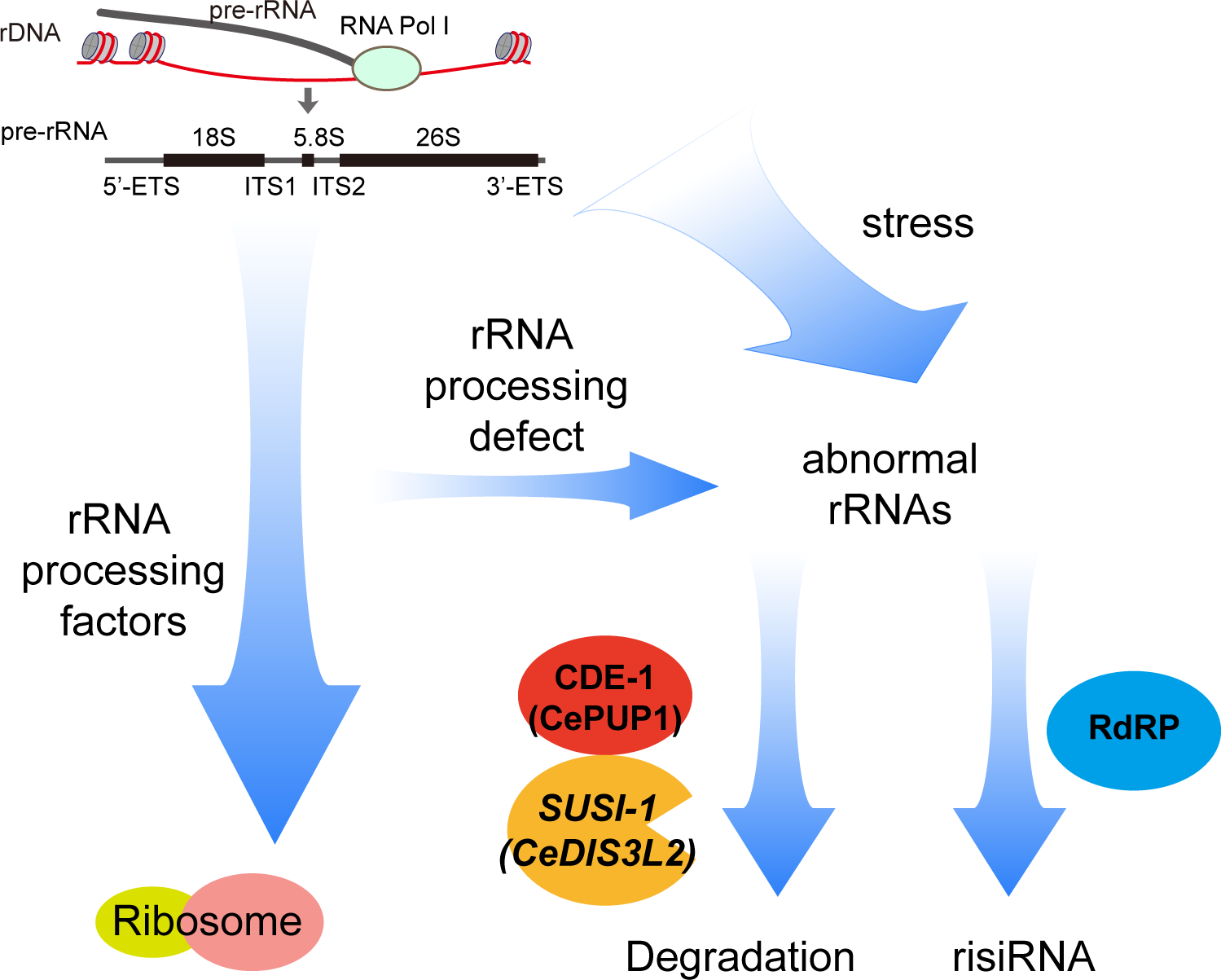
A working model of risiRNA biogenesis in *C. elegans*. The erroneous cellular rRNAs are scrutinized and suppressed through a number of mechanisms. Erroneous rRNAs are uridylated by CDE-1 and degraded by exoribonucleases such as SUSI-1(ceDis3L2). The disruption of CDE-1 or SUSI-1(ceDis3L2) results in the accumulation of erroneous rRNAs that thereafter recruit RdRPs to synthesize risiRNA. A risiRNA-mediated feedback loop silences rRNA expression through RNAi machinery and compensates for the disruption of the degradation of erroneous rRNA transcripts.

3’-end modifications play important roles in regulating the stability of siRNAs via distinct mechanisms as well. For example, methylation of the 3’-end inhibits uridylation and correlates with increased steady state levels of small RNAs (Kamminga, Luteijn et al., 2010, Ren, Xie et al., 2014). In contrary, 3’ terminal uridylation may promote the degradation of siRNA (Ibrahim, Rymarquis et al., 2010, van Wolfswinkel et al., 2009). Among the three PUP proteins in *C. elegans*, CDE-1 uridylates endogenous siRNAs and modulates their binding affinity to CSR-1 and WAGO-4 (van Wolfswinkel et al., 2009, Xu et al., 2018). PUP-2 has been reported to target *let-7* miRNA (Heo et al., 2012, Lehrbach et al., 2009). Although PUP-3 has been validated as uridyl transferase, its targets are still unclear. Here, we found that CDE-1, but not PUP-2 or PUP-3, are engaged in suppressing risiRNA production. How these PUPs recognize their specific targets is an enigma.

Additional questions remain as to how and why erroneous rRNAs could be recognized by CDE-1. Our previous work found that either the modification errors or processing errors of rRNAs trigger the generation of risiRNAs (Zhu et al., 2018). How these different kinds of errors are sensed and scrutinized is still unknown. Deciphering the intricate interaction network of CDE-1 or other TUTases is key to fully understanding the effect of RNA uridylation. In addition, CDE-1 was previously reported required for RNAi inheritance (Spracklin et al., 2017, Xu et al., 2018). The underlying mechanism remains unclear. Here, we found that CDE-1 interacted with SUSI-1, another protein required for the inheritance of RNAi. Further elucidating the function of SUSI-1 and CDE-1 will shed light on the mechanism of RNAi inheritance.

## Materials and methods

### Strains

Bistol strain N2 was used as the standard wild-type strain. All strains were grown at 20°C unless otherwise specified. The strains used in this study were listed in supplementary Table S1.

### Quantification of the subcellular location of NRDE-3

The subcellular localization of NRDE-3 was quantified as described previously (Zhou et al., 2017b). Images were collected on a Leica DM4B microscope.

### Quantitative RT-PCR

RNA was isolated from the indicated animals and subjected to DNase I digestion (Thermo Fisher). cDNA was generated from the isolated RNA using a **GoScript*™ Reverse_Transcription_System* (Promega) according to the vendor’s protocol. qPCR was performed using a MyIQ2 real-time PCR system (Bio-Rad) with an AceQ SYBR Green Master mix (Vazyme). The primers used in RT-qPCR were listed in Table S2. *eft-*3 mRNA was used as an internal control for sample normalization. Data analysis was performed using a comparative threshold cycle (ΔΔCT) approach.

### Brood size

Synchronized L3 worms were individually placed onto fresh NGM plates, and the progeny numbers were scored.

### Construction of plasmids and transgenic strains

For *CDE-1::GFP*, a *cde-1* promoter and CDS region were PCR-amplified with the primers 5′-TACGACTCACTAGTGGGCAGgacgtgggacataaacgaagaaag-3′ and 5′-ATAGCTCCACCTCCACCTCCTTTGTTGTACGAGCGATGATAG-3′ from N2 genomic DNA. A *GFP::3×FLAG* region was PCR-amplified with the primers 5′-GGAGGTGGAGGTGGAGCTATGAGTAAAGGAGAAGAAC-3′ and 5′-TCACTTGTCATCGTCATCCT-3′ from plasmid pSG085. The CDE-1 3′ UTR (untranslated region) was PCR-amplified with the primers 5′-ACAAGGATGACGATGACAAGTAAattctctccacccattcac-3′ and 5′-CTACGTAATACGACTCACTTaactgatcggttgcttctctcac-3′ from N2 genomic DNA. A ClonExpress MultiS One-step Cloning Kit (Vazyme, C113-02) was used to insert the *CDE-1::GFP::3×FLAG* fusion gene into the pCFJ151 vector. The transgene was integrated into *C. elegans* chromosome II by the MosSCI method (Frokjaer-Jensen, Davis et al., 2014). Using the same method, the *CDE-1::mCherry* fusion gene was integrated into *C. elegans* chromosome Ⅴ.

The primers used for dual-sgRNA-directed CRISPR/Cas9-mediated *cde-1* gene deletion were 5′-TCCGGATAGTGATTACAATG-3′ and 5′-GGTATTATGTTGAACGACAT-3′.

For *3×FLAG::GFP::SUSI-1*, the predicted *susi-1* promoter was PCR-amplified with the primers 5′-TACGACTCACTAGTGGGCAGtatcagggagattctgctgtg-3′ and 5′-tcatggtctttgtagtccatACTTTCAACTGCTGACATctag-3′ from N2 genomic DNA. The *3×FLAG::GFP* coding region was PCR amplified from plasmid pSG085 with the primers 5′-AGCTCTTCCTATGGACTACAAAGACCATGAC-3′ and 5′-ATAGCTCCACCTCCACCTCCTTTGTATAGTTCATCCATGCC-3′. The SUSI-1 coding region and the predicted 3′ UTR were then amplified by PCR from N2 genomic DNA with primers 5′-AAGGAGGTGGAGGTGGAGCTATGTCAGCAGTTGAAAGTCCCG-3′ and 5′-CTACGTAATACGACTCACTTGTGTGGATTAACACAGCCAATTG-3′ from N2 genomic DNA. The ClonExpress MultiS One-step Cloning Kit (Vazyme, C113-02) was used to insert the *3×FLAG::GFP::SUSI-1* fusion gene into the pCFJ151 vector. The transgene was integrated into *C. elegans* chromosome II by the MosSCI system.

### RNA immunoprecipitation (RIP)

RIP experiments were performed as previously described with hypochlorite-isolated embryos of indicated animals (Zhou et al., 2017b). The embryos were sonicated in lysis buffer (20 mM Tris-HCl (pH 7.5), 200 mM NaCl, 2.5 mM MgCl_2_, and 0.5% NP-40), precleared with protein G-agarose beads (Roche), and incubated with anti-FLAG M2 agarose beads (Sigma #A2220). The beads were washed extensively, and 3*×*FLAG::GFP-tagged protein and associated RNAs were eluted with 100 μg/mL 3*×*FLAG peptide (Sigma). The eluates were incubated with TRIzol reagent (Invitrogen), which was followed by isopropanol precipitation. Then, small RNAs were quantified by deep sequencing.

### Deep sequencing of small RNAs and bioinformatic analysis

Total RNAs and the Argonaute-associated RNAs were isolated from the indicated animals and subjected to small RNA deep sequencing using an Illumina platform (Novogene Bioinformatics Technology Co., Ltd.), as previously described (Zhou et al., 2017b).

For Argonaute-associated RNAs, synchronized worms were sonicated in sonication buffer (20 mM Tris-HCl,pH 7.5, 200 mM NaCl, 2.5 mM MgCl_2_, and 0.5% NP40). The eluates were incubated with TRIzol reagent and then precipitated with isopropanol. The precipitated RNA was treated with FastAP Thermosensitive Alkaline Phosphatase (Thermo Scientific), re-extracted with TRIzol, and treated with T4 Polynucleotide Kinase (T4 PNK, Thermo Scientific) in the presence of 1 mM ATP before library construction.

Small RNAs were subjected to deep sequencing using an Illumina platform (Novogene Bioinformatics Technology Co., Ltd.). Briefly, small RNAs ranging from 18 to 30 nt were gel-purified and ligated to a 3’ adaptor (5’-pUCGUAUGCCGUCUUCUGCUUGidT-3’; p, phosphate; idT, inverted deoxythymidine) and a 5’ adaptor (5’-GUUCAGAGUUCUACAGUCCGACGAUC-3’), respectively. The ligation products were gel-purified, reverse transcribed, and amplified using an Illumina sRNA primer set (5’-CAAGCAGAAGACGGCATACGA-3’; 5’-AATGATACGGCGACCACCGA-3’). The samples were then sequenced using an Illumina HiSeq platform.

The Illumina-generated raw reads were first filtered to remove adaptors, low-quality tags and contaminants to obtain clean reads by Novogene. Clean reads ranging from 18 to 30 nt were mapped to the transcriptome assembly WS243 using Bowtie2 with default parameters. The number of reads targeting each transcript were counted by custom Perl scripts. The number of total reads mapped to the genome minus the number of total reads corresponded to sense rRNA transcripts (5S, 5.8S, 18S and 26S), which was used as the normalization number, to exclude the possible degradation fragments of sense rRNAs.

### Proteomic analysis

Proteomic analysis was conducted as previously described (Zeng, Weng et al., 2019). Briefly, mixed-stage transgenic worms expressing CDE-1::GFP were resuspended in equal volumes of 2*×* lysis buffer (50 mM Tris-HCl pH 8.0, 300 mM NaCl, 10% glycerol, 1% Triton X-100, Roche®cOmplete EDTA-free Protease Inhibitor Cocktail, 10 mM NaF, and 2 mM Na_3_VO_4_), and lysed in a FastPrep-24 5G homogenizer. The lysate supernatant was incubated with anti-GFP antibody, which was linked to beads, for one hour at 4 ଌ. The beads were then washed three times with cold lysis buffer. The GFP immunoprecipitates were eluted with chilled elution buffer (100 mM glycine-HCl, pH 2.5). Approximately 1/8 of the eluates were subjected to western blotting analysis. The rest of the eluates were precipitated with TCA or cold acetone and dissolved in 100 mM Tris (pH 8.5), with 8 M urea. The proteins were reduced with TCEP, alkylated with IAA, and finally digested with trypsin at 37 ଌ overnight. LC-MS/MS analysis of the resulting peptides and MS data processing approaches were conducted as previously described (Feng, Zhu et al., 2017). A WD scoring matrix was used to identify high-confidence candidate interacting proteins.

### Coimmunoprecipitation analysis

The lysates of transgenic worms were prepared using RIP lysis buffer (50 mM Tris (pH 7.4), 150 mM NaCl, 1% NP-40, 0.1% SDS, 1 mM EDTA, 0.5% sodium deoxycholate, and protease inhibitors (Thermo)]. Immunoprecipitations with anti-FLAG® M2 affinity gel (a2220, Sigma) or agarose beads (ab193255, Abcam) with anti-GFP antibody (ab290, Abcam) and anti-SUSI-1 antibody (lot number 20121105, Abmart) were performed at 4 °C overnight. Protein complexes were eluted by boiling in 2× SDS loading buffer. Anti-GFP, anti-SUSI-1 and anti-Actin (Servicebio GB12001) antibodies that were used for western blots were diluted to 1:2000, 1:500 and 1:5000, respectively.

### rRNA 3’ tail-seq

26S rRNA tail-seq was conducted as described previously (Zhou et al., 2017b). Briefly, total RNA were extracted from embryos or L3 larva, digested by DNase I, and then ligated to the following 3′ RNA linkers with T4 RNA ligase (Thermo #EL0021) (1 μg total RNA, 2 μl 3′ RNA linker (10 μM), 1 μl 10× T4 RNA ligation buffer, 2 μl T4 RNA ligase) by incubating at 37 °C for 30 min.

3′ RNA linker: 5′-p<u>GATCC</u>ACACTCGGGCACCAAGGATTTAACCGCGAATTCCAGC-NH2-3′ (the underlined sequence served as a barcode for sample labeling).

The RNAs were reverse transcribed with the following primers: 3′ linker RT: 5′-GCTGGAATTCGCGGTTAAATCCTTGGTGCCCGAGTGT<u>GGATC</u>-3′. The cDNAs were PCR amplified with the primers 26S rRNA-F: 5′-CAGATCACTCTGGTTCAATGTC-3′ and 3′ linker RT primers, gel purified and then deep sequenced using an Illumina platform, according to the manufacturer′s instructions, by Novogene (Beijing, China). The number of reads with distinct 3′ -end modifications were counted by custom Perl scripts.

### RNAi inheritance assay

Synchronized L1 animals of the indicated genotypes were exposed to bacteria expressing *gfp* dsRNA. F1 embryos were collected by hypochlorite/NaOH treatment and grown on HT115 control bacteria. The GFP expression levels in both the parental generation and the progeny were visualized and scored. Images were collected with a Leica DM4B microscope system.

### Statistics

Bar graphs with error bars represent the mean and SD. All of the experiments were conducted with independent *C. elegans* animals for the indicated N replicates. Statistical analysis was performed with two-tailed Student’s t-tests or unpaired Wilcoxon tests. The threshold for Student’s t-test *p* values was set to 0.05.

## Acknowledgments

We are grateful to Drs. Shanhui Liao and Shuai Wei, and the members of the Guang lab for their comments. We are grateful to the International *C. elegans* Gene Knockout Consortium and the National Bioresource Project for providing the strains. Some strains used in this study were provided by the CGC, which is funded by NIH Office of Research Infrastructure Programs (P40 OD010440). This work was supported by grants from the China Postdoctoral Science Foundation (2015M582006), the Natural Science Foundation of Anhui Province (1608085MC68), and the National Natural Science Foundation of China (Nos. 31671346, 91640110, 31870812 and 31871300). This study was supported, in part, by Hefei National Science Center Pilot Project Funds and CAS Interdisciplinary Innovation Team.

## Author Contributions

Y.W., C.W., C. Z., and S.G. designed the experiments. Y.W., C.W., X.C., and X.H. performed experiments. Y.W. and C.W. analyzed the data. Y.W., C.W., and S.G. wrote the manuscript. All authors have discussed the manuscript.

## Declaration of Interests

The authors declare no competing financial interests.

---

**Figure S1.**
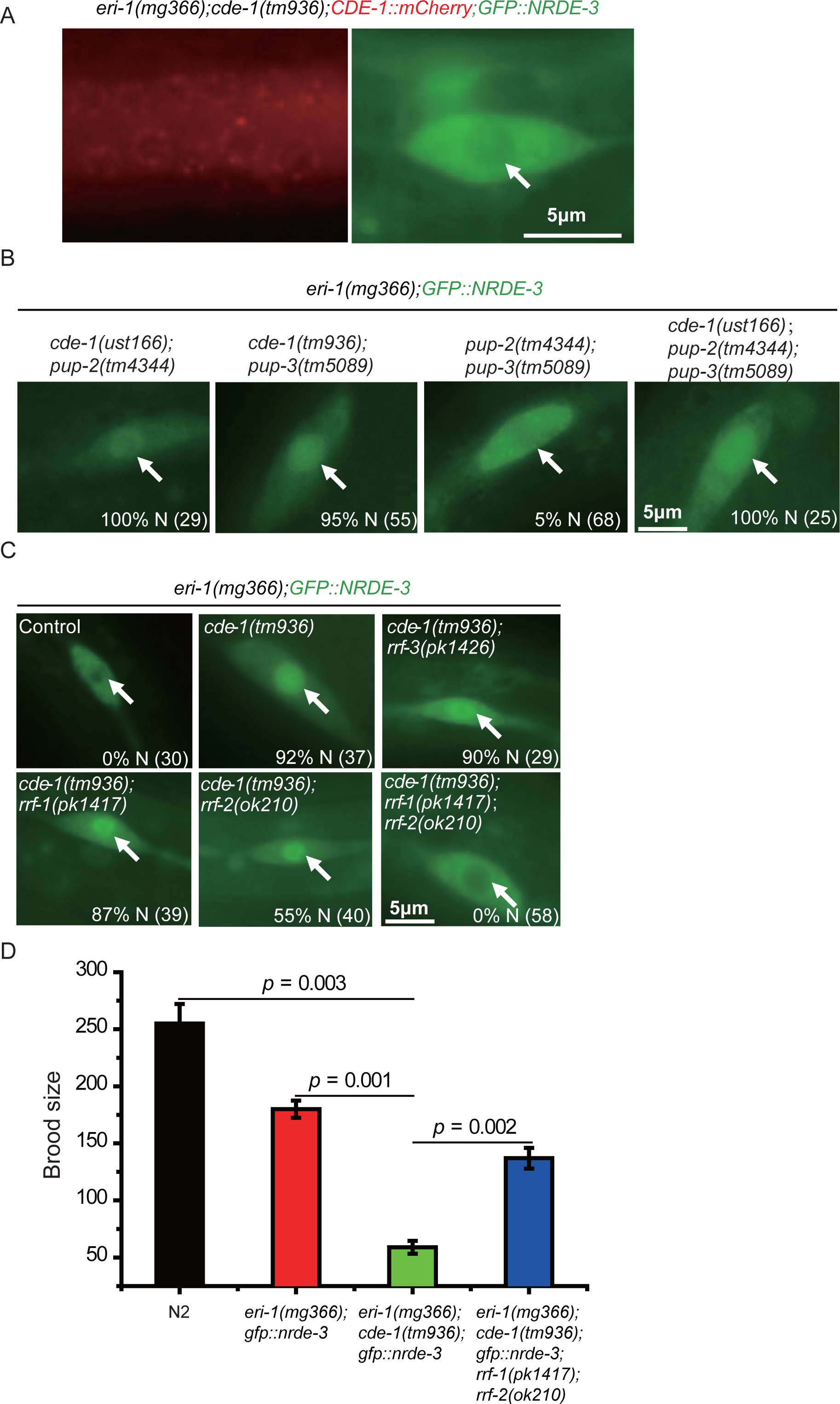
*rrf-1* and *rrf-2* were required for risiRNA production in the *cde-1* mutant. (A) *CDE-1::mCherry* was able to redistribute NRDE-3 from the nucleus to the cytoplasm in *cde-1(tm936)* mutants. Indicates were the seam cells of indicated animals. White arrows, nucleus. (B) The depletion of *pup-2* and *pup-3* together was not able to redistribute NRDE-3 from the cytoplasm to the nucleus. Indicates were the seam cells of indicated animals. The numbers indicated the percentage of animals with nuclear enriched NRDE-3 in seam cells. The number of scored animals is indicated in the parentheses. White arrows, nucleus. (C) *rrf-1* and *rrf-2* were required for risiRNA production. Images are of representative seam cells. (D) The depletion of *rrf-1* and *rrf-2* partially restored the fecundity of *eri-1(mg366);cde-1(tm936)* animals. Data are presented as the mean ± s.d. n = 3.

**Figure S2.**
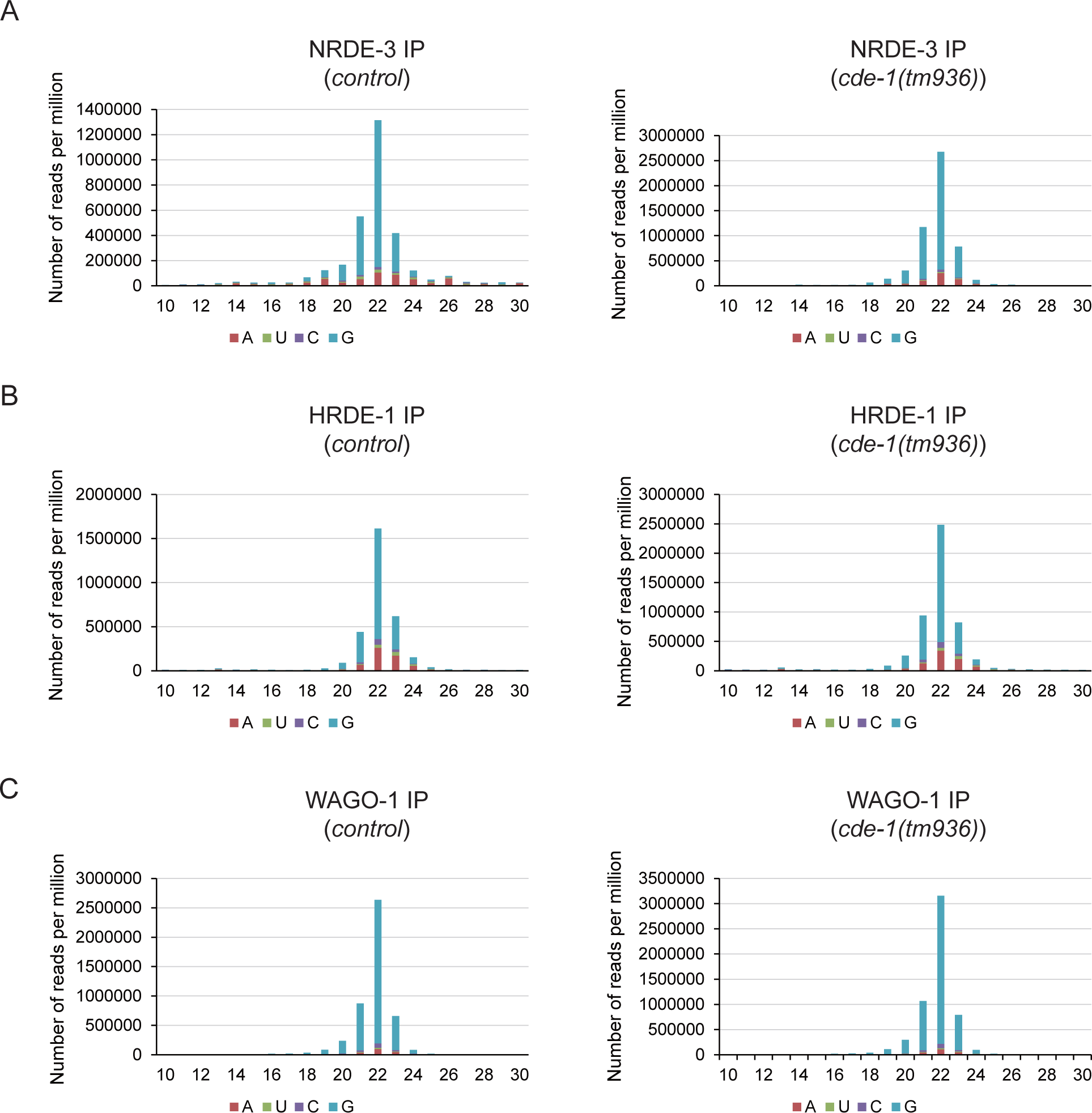
Size distribution and 5’ end nucleotide preference of siRNAs identified by deep sequencing. (A) NRDE-3-, (B) HRDE-1-, and (C) WAGO-1-bound small RNAs in indicated animals were deep sequenced. Size distribution and 5’ end nucleotide preference were analyzed.

**Figure S3.**
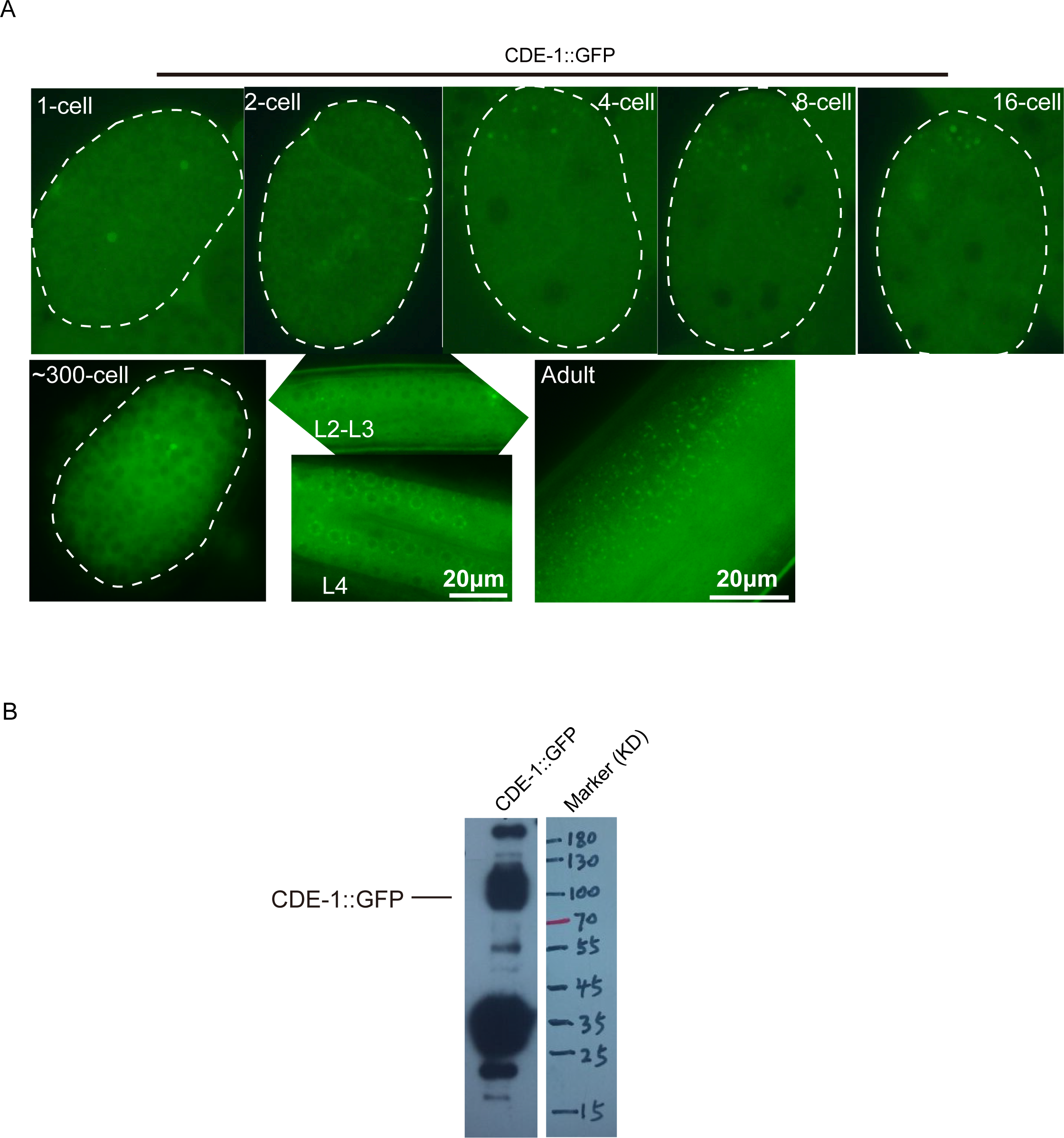
The expression pattern of CDE-1 at indicated developmental stages. (A) CDE-1::GFP was visualized by fluorescent microscopy at indicated developmental stages. (B) Western blotting analysis of CDE-1::GFP was performed after GFP immunoprecipitation.

**Table S1:**
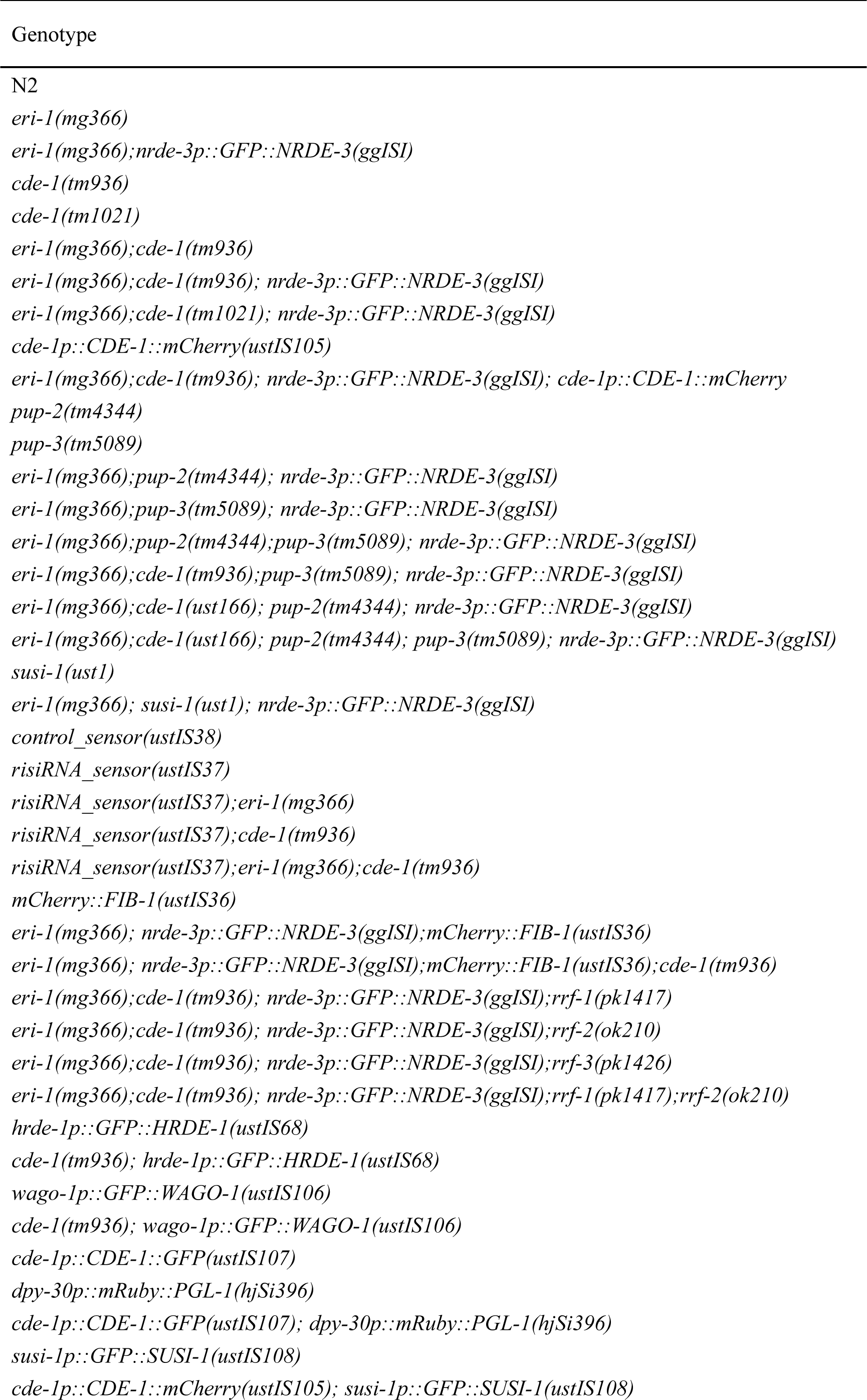
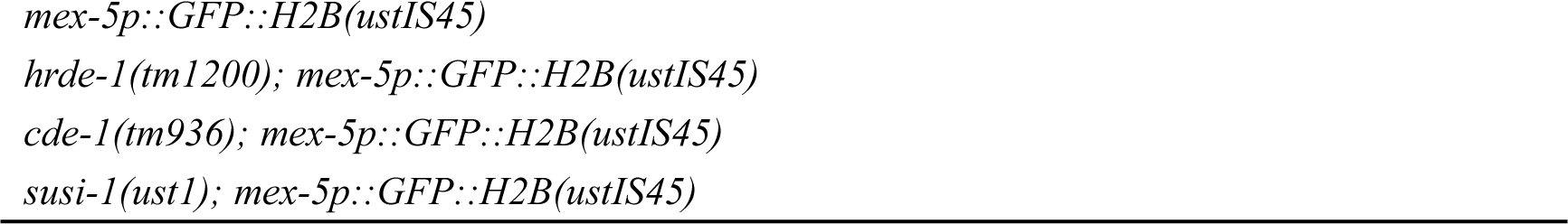
Strains used in this work.

**Table S2:**
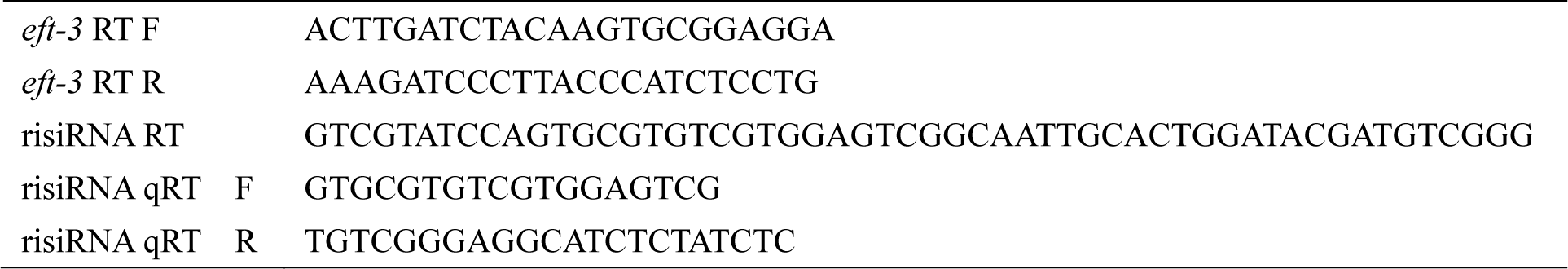
Primers used for quantitative real-time PCR analysis.

## References

Blahna MT, Jones MR, Quinton LJ, Matsuura KY, Mizgerd JP (2011) Terminal uridyltransferase enzyme Zcchc11 promotes cell proliferation independent of its uridyltransferase activity. J Biol Chem 286: 42381–9

Claycomb JM, Batista PJ, Pang KM, Gu W, Vasale JJ, van Wolfswinkel JC, Chaves DA, Shirayama M, Mitani S, Ketting RF, Conte D, Jr., Mello CC (2009) The Argonaute CSR-1 and its 22G-RNA cofactors are required for holocentric chromosome segregation. Cell 139: 123–34

De Almeida C, Scheer H, Zuber H, Gagliardi D (2018) RNA uridylation: a key posttranscriptional modification shaping the coding and noncoding transcriptome. Wiley Interdisciplinary Reviews-Rna 9

Faehnle CR, Walleshauser J, Joshua-Tor L (2014) Mechanism of Dis3l2 substrate recognition in the Lin28-let-7 pathway. Nature 514: 252–256

Feng GX, Zhu ZW, Li WJ, Lin QR, Chai YP, Dong MQ, Ou GS (2017) Hippo kinases maintain polarity during directional cell migration in Caenorhabditis elegans. Embo Journal 36: 334–345

Frokjaer-Jensen C, Davis MW, Sarov M, Taylor J, Flibotte S, LaBella M, Pozniakovsky A, Moerman DG, Jorgensen EM (2014) Random and targeted transgene insertion in Caenorhabditis elegans using a modified Mos1 transposon. Nat Methods 11: 529–34

Guang S, Bochner AF, Pavelec DM, Burkhart KB, Harding S, Lachowiec J, Kennedy S (2008) An Argonaute transports siRNAs from the cytoplasm to the nucleus. Science 321: 537–41

Hagan JP, Piskounova E, Gregory RI (2009) Lin28 recruits the TUTase Zcchc11 to inhibit let-7 maturation in mouse embryonic stem cells. Nat Struct Mol Biol 16: 1021–5

Henras AK, Plisson-Chastang C, O’Donohue MF, Chakraborty A, Gleizes PE (2015) An overview of preribosomal RNA processing in eukaryotes. Wiley Interdiscip Rev RNA 6: 225–42

Heo I, Ha M, Lim J, Yoon MJ, Park JE, Kwon SC, Chang H, Kim VN (2012) Mono-uridylation of pre-microRNA as a key step in the biogenesis of group II let-7 microRNAs. Cell 151: 521–32

Houseley J, LaCava J, Tollervey D (2006) RNA-quality control by the exosome. Nature Reviews Molecular Cell Biology 7: 529–539

Ibrahim F, Rymarquis LA, Kim EJ, Becker J, Balassa E, Green PJ, Cerutti H (2010) Uridylation of mature miRNAs and siRNAs by the MUT68 nucleotidyltransferase promotes their degradation in Chlamydomonas. Proc Natl Acad Sci U S A 107: 3906–11

Jones MR, Quinton LJ, Blahna MT, Neilson JR, Fu S, Ivanov AR, Wolf DA, Mizgerd JP (2009) Zcchc11-dependent uridylation of microRNA directs cytokine expression. Nat Cell Biol 11: 1157–63

Kamminga LM, Luteijn MJ, den Broeder MJ, Redl S, Kaaij LJ, Roovers EF, Ladurner P, Berezikov E, Ketting RF (2010) Hen1 is required for oocyte development and piRNA stability in zebrafish. EMBO J 29: 3688–700

Karbstein K (2013) Quality control mechanisms during ribosome maturation. Trends Cell Biol 23: 242–50

Kwak JE, Wickens M (2007) A family of poly(U) polymerases. RNA 13: 860–7

Lafontaine DL (2010) A ‘garbage can’ for ribosomes: how eukaryotes degrade their ribosomes. Trends Biochem Sci 35: 267–77

Lafontaine DLJ (2015) Noncoding RNAs in eukaryotic ribosome biogenesis and function. Nature Structural & Molecular Biology 22: 11–19

Le Pen J, Jiang H, Di Domenico T, Kneuss E, Kosalka J, Leung C, Morgan M, Much C, Rudolph KLM, Enright AJ, O’Carroll D, Wang D, Miska EA (2018) Terminal uridylyltransferases target RNA viruses as part of the innate immune system. Nat Struct Mol Biol 25: 778–786

Lee LW, Lee CC, Huang CR, Lo SJ (2012) The nucleolus of Caenorhabditis elegans. J Biomed Biotechnol 2012: 601274

Lee M, Kim B, Kim VN (2014) Emerging roles of RNA modification: m(6)A and U-tail. Cell 158: 980–987

Lehrbach NJ, Armisen J, Lightfoot HL, Murfitt KJ, Bugaut A, Balasubramanian S, Miska EA (2009) LIN-28 and the poly(U) polymerase PUP-2 regulate let-7 microRNA processing in Caenorhabditis elegans. Nat Struct Mol Biol 16: 1016–20

Li Y, Maine EM (2018) The balance of poly(U) polymerase activity ensures germline identity, survival and development in Caenorhabditis elegans. Development 145

Lubas M, Damgaard CK, Tomecki R, Cysewski D, Jensen TH, Dziembowski A (2013) Exonuclease hDIS3L2 specifies an exosome-independent 3’-5’ degradation pathway of human cytoplasmic mRNA. Embo Journal 32: 1855–1868

Menezes MR, Balzeau J, Hagan JP (2018) 3’ RNA Uridylation in Epitranscriptomics, Gene Regulation, and Disease. Frontiers in Molecular Biosciences 5

Pena C, Hurt E, Panse VG (2017) Eukaryotic ribosome assembly, transport and quality control. Nat Struct Mol Biol 24: 689–699

Pirouz M, Du P, Munafo M, Gregory RI (2016) Dis3l2-Mediated Decay Is a Quality Control Pathway for Noncoding RNAs. Cell Rep 16: 1861–73

Pirouz M, Munafo M, Ebrahimi AG, Choe J, Gregory RI (2019) Exonuclease requirements for mammalian ribosomal RNA biogenesis and surveillance. Nat Struct Mol Biol 26: 490–500

Ren G, Xie M, Zhang S, Vinovskis C, Chen X, Yu B (2014) Methylation protects microRNAs from an AGO1-associated activity that uridylates 5’ RNA fragments generated by AGO1 cleavage. Proc Natl Acad Sci U S A 111: 6365–70

Schmidt K, Butler JS (2013) Nuclear RNA surveillance: role of TRAMP in controlling exosome specificity. Wiley Interdiscip Rev RNA 4: 217–31

Spracklin G, Fields B, Wan G, Becker D, Wallig A, Shukla A, Kennedy S (2017) The RNAi Inheritance Machinery of Caenorhabditis elegans. Genetics 206: 1403–1416

Thoms M, Thomson E, Bassler J, Gnadig M, Griesel S, Hurt E (2015) The Exosome Is Recruited to RNA Substrates through Specific Adaptor Proteins. Cell 162: 1029–1038

Trippe R, Guschina E, Hossbach M, Urlaub H, Luhrmann R, Benecke BJ (2006) Identification, cloning, and functional analysis of the human U6 snRNA-specific terminal uridylyl transferase. Rna-a Publication of the Rna Society 12: 1494–1504

Ustianenko D, Pasulka J, Feketova Z, Bednarik L, Zigackova D, Fortova A, Zavolan M, Vanacova S (2016) TUT-DIS3L2 is a mammalian surveillance pathway for aberrant structured non-coding RNAs. EMBO J 35: 2179–2191

van Wolfswinkel JC, Claycomb JM, Batista PJ, Mello CC, Berezikov E, Ketting RF (2009) CDE-1 affects chromosome segregation through uridylation of CSR-1-bound siRNAs. Cell 139: 135–48

Vanacova S, Stefl R (2007) The exosome and RNA quality control in the nucleus. EMBO Rep 8: 651–7

Warkocki Z, Krawczyk PS, Adamska D, Bijata K, Garcia-Perez JL, Dziembowski A (2018) Uridylation by TUT4/7 Restricts Retrotransposition of Human LINE-1s. Cell 174: 1537-+

Xu F, Feng XZ, Chen XY, Weng CC, Yan Q, Xu T, Hong MJ, Guang SH (2018) A Cytoplasmic Argonaute Protein Promotes the Inheritance of RNAi. Cell Reports 23: 2482–2494

Yan Q, Zhu C, Guang S, Feng X (2019) The Functions of Non-coding RNAs in rRNA Regulation. Front Genet 10: 290

Yeo J, Kim VN (2018) U-tail as a guardian against invading RNAs. Nature Structural & Molecular Biology 25: 903–905

Yi YH, Ma TH, Lee LW, Chiou PT, Chen PH, Lee CM, Chu YD, Yu H, Hsiung KC, Tsai YT, Lee CC, Chang YS, Chan SP, Tan BC, Lo SJ (2015) A Genetic Cascade of let-7-ncl-1-fib-1 Modulates Nucleolar Size and rRNA Pool in Caenorhabditis elegans. PLoS Genet 11: e1005580

Zeng C, Weng C, Wang X, Yan YH, Li WJ, Xu D, Hong M, Liao S, Dong MQ, Feng X, Xu C, Guang S (2019) Functional Proteomics Identifies a PICS Complex Required for piRNA Maturation and Chromosome Segregation. Cell Rep 27: 3561–3572 e3

Zhou X, Chen X, Wang Y, Feng X, Guang S (2017a) A new layer of rRNA regulation by small interference RNAs and the nuclear RNAi pathway. RNA Biol 14: 1492–1498

Zhou XF, Feng XZ, Mao H, Li M, Xu F, Hu K, Guang SH (2017b) RdRP-synthesized antisense ribosomal siRNAs silence pre-rRNA via the nuclear RNAi pathway. Nature Structural & Molecular Biology 24: 258-+

Zhu C, Yan Q, Weng C, Hou X, Mao H, Liu D, Feng X, Guang S (2018) Erroneous ribosomal RNAs promote the generation of antisense ribosomal siRNA. Proc Natl Acad Sci U S A 115: 10082–10087

